# A novel method revealing animal evolutionary relationships based on milk Mid-infrared

**DOI:** 10.1101/2022.11.18.517067

**Authors:** Yikai Fan, Jiazheng Han, Haitong Wang, Liangkang Nan, Xuelu Luo, Chu Chu, Liang Wang, Li Liu, Yongqing Li, Chunfang Li, Xiaoli Ren, Lei Ding, Wenju Liu, Xingjie Hao, Yansen Chen, George E. Liu, Yang Zhou, Shujun Zhang

**Author notes:** These authors contributed equally: Yikai Fan, Jiazheng Han, Haitong Wang.

## Abstract

Mid-infrared spectra (MIRS) can effectively reflect the chemical bonds in milk, which has been widely used in dairy herd improvement. However, the relationship between MIRS and animal evolution remains largely unclear. This study firstly found great differences in MIRS and the components of milk by analyzing MIRS information of 12 different mammal species. A five-level discriminant model of evolutionary level based on MIRS was established with a test set kappa coefficient >0.97. In addition, a regression model of genetic distance was also established to estimate the genetic distance of different animal species with a correlation coefficient of R >0.94. These results showed that this method could be used for accurate mammalian evolutionary relationship assessment. We further clarified the potential relationship between MIRS and genes, such as PPP3CA and SCD that could change MIRS by regulating specific milk components. In conclusion, we expand the application of MIRS in animal species identification and evolution research and provide new perspectives for the research on the formation mechanism of different animal milk special components.

## Introduction

To meet the need for timely and rapid detection of animal milk and dairy products, infrared spectrum (IRS) was used in 1967 to measure the fat content (FP, fat percentage), protein content (PP, protein percentage), and lactose content (LP, lactose percentage)^[1]^. Mid-infrared spectra (MIRS) are the fundamental frequency absorption band of organic matter and inorganic ions. The position, shape, number, and intensity of MIRS characteristic peaks of animal milk can reflect the components and content of milk, and thus it is especially suitable for liquid milk analysis^[2,3]^. MIRS detection technology has been widely used in milk component detection and global dairy herd improvement (DHI) due to its multiple advantages of low cost, simplicity, rapidity, batch, accuracy, and non-destructiveness.

Different species of mammals (referred to as animals) exhibit great differences in the content of milk components. The Perissodactyla horse and ass milk have low FP (1.21%) and high LP (6.37%) ^[4–6]^. The milk of Artiodactyla camel has lower FP and PP (6% and 4%) than that of pigs (9.97% and 5.25%), while the PP of pig colostrum is as high as 17%^[7, 8]^. FP and PP (5.3-9.3% and 4.5-6.6%) of goat and sheep milk are higher than those of cows ^[9,10]^. FP and PP (8.5% and 4.5%) of buffalo milk are higher than those of Holstein cattle with a high fatty acid content^[11,12]^. The contents of FP, PP, and calcium in yak milk are 1.68-2 times as high as those in Holstein milk^[13]^. PP (3.32%-3.61%) in Jersey and Simmental milk is higher than that in Holstein, whereas FP and PP in Holstein milk are not less than 3.4% and 2.9%^[14, 15]^.

There are also great differences in MIRS among different animals. The CH_2_ absorption bands of goat milk, sheep milk, and cow milk are different at 2927cm^-1^, and the MIRS absorption peak rises with the increasing FP. The MIRS absorption peaks associated with PP (amide I near 1654cm^-1^ and amide II near 1544cm^-1^) and LP (1159 cm^-1^ and 1076 cm^-1^) of sheep milk are higher than those of cow milk and goat milk^[16]^. At present, only the research on the MIRS bands corresponding to the main milk components has been carried out, but no reports on more characteristic MIRS bands and their application have been available.

MIRS wave points are hereditary. The heritability of 1060 wave points of MIRS is 0.003-0.42,which is similar to the estimated value of milk-related traits, indicating that milk MIRS is as heritable as milk components^[17]^. The heritability of MIRS wave points can reach the medium level and above (0-0.63), and thus it could be speculated that genes affect MIRS wave points by regulating milk components^[18]^.The heritability of MIR wave points in Holstein and Jersey ranges from 0 to 0.31, and MIR wave points and genomes are significantly correlated^[19]^. Our previous study has shown that the heritability of 1060 MIRS wave points is 0.04 on average, which is similar to the heritability of FP, PP, and LP^[20]^.

MIRS-based classification or regression models can be established by MIRS feature selection, training, testing, and primary model optimization based on machine learning. The MIRS features can be selected using principal component analysis (PCA) and uninformative variable elimination (UVE). In addition, random forest (RF), support vector machines (SVM), partial least squares regression (PLSR), and artificial neural network (ANN) are employed to establish classification and regression models, and the optimal model is selected according to accuracy and kappa coefficient.

Animal evolutionary relationships are mainly established by such algorithms as the neighbor-joining method (NJ) or Bayes method based on the differences in genome nucleic acid sequences^[21,22]^. With the gradual implementation of *The Earth BioGenome Project* (EBP), the genetic maps and evolutionary relationships of a large number of organisms have been published^[23]^. Although the method of constructing phylogenetic trees based on nucleic acid sequence is accurate, the cost of whole genome sequencing is high, and thus it is necessary to develop a method that can replace genetic material to construct the evolutionary relationship of animals.

To sum up, there are differences in the contents of milk components and MIRS among different animals. MIRS can reflect the characteristics of milk components, and it is hereditary. Based on this, it could be speculated that it is feasible to use MIRS to construct animal evolutionary relationships. At present, the genome is mainly used to analyze animal evolution, and there have been few literature reports on animal evolution research based on MIRS. MIRS technology has multiple advantages such as inexpensive, rapidity, batch, accuracy, simplicity, and non-destructiveness. Therefore, the main purpose of this study is to (1) construct the evolutionary relationship of animals by using MIRS; (2) identify the evolution-related milk components and their characteristic wavebands, and analyze their change regularity in the evolution process; (3) reveal the relationship between characteristic wavebands and genes to resolve the functions and genetic information of MIR characteristic spectra. The significance of the research lies in (1) breaking through the traditional technology dependent on genetic materials to clarify the evolutionary relationship of animals, and creating a new technology for establishing an animal evolutionary relationship based on MIRS; (2) revealing the formation mechanism of different animal milk-specific or highly expressed components, and providing new methods and perspectives for the research on the evolution processes and characteristics of different animal species.

## Results

### 2.1 Great differences in milk components and MIRS among different species of mammals

#### 2.1.1 Differences in milk components among different animals

The contents of 23 milk components in 12 animal species (Table 2, except OBY with small sample size) were compared. As shown in Table S2, horses and asses exhibited the highest lactose percentage (LP, 7.04%, 6.51%), and lowest fat percentage (FP, 0.54%, 0.35%), protein percentage (PP, 1.38%, 1.35%), and total solid contents (TS, 9.81%, 8.84%). Buffalo had the highest FP (7.66%) and a relatively higher level of other milk components. Goat and camel milk displayed moderate milk component contents. Yak showed relatively high contents of FP (6.45%) and TS (17.03%) and low content of fatty acids. Taken together, the 12 animal milk components were quite different in their own characteristics.

**Table 1.** Characteristic wavebands of different milk components.

| Milk components | MIRS characteristic band |
| --- | --- |
| Fat | 1180.55-1257.71cm<sup>-1</sup> 1728.38-1770.82cm<sup>-1</sup> 2839.49-2966.80cm<sup>-1</sup> |
| Lactose | 1014.65-1134.25cm<sup>-1</sup> 1311.72-1373.45cm<sup>-1</sup> |
| Protein & Total-CN | 1396.60-1415.89cm<sup>-1</sup> 1431.32cm<sup>-1</sup> 1473.76cm<sup>-1</sup> 1405.62-1585.64cm<sup>-1</sup> |
| TS | 1219.13-1273.14cm<sup>-1</sup> |
| κ-CN | 1435.18-1473.76cm<sup>-1</sup> 1504.62-1581.78cm<sup>-1</sup> |
| β-CN | 1219.13-1265.42cm<sup>-1</sup> 1446.75-1570.21cm<sup>-1</sup> |
| αS1-CN | 3695.96-3703.68cm<sup>-1</sup> |
| α-LA | 925.92-1006.94cm<sup>-1</sup> |
| Total_AA | 1709.09-1720.67cm<sup>-1</sup> |
| Arg | 1477.614-1593.354cm<sup>-1</sup> |
| TAU | 3533.93cm<sup>-1</sup> |
| PETH | 1292.43-1346.44cm<sup>-1</sup> |
| FA | 1427.46-1481.47cm<sup>-1</sup> 3067.11-3155.84cm<sup>-1</sup> |

**Table 2.**
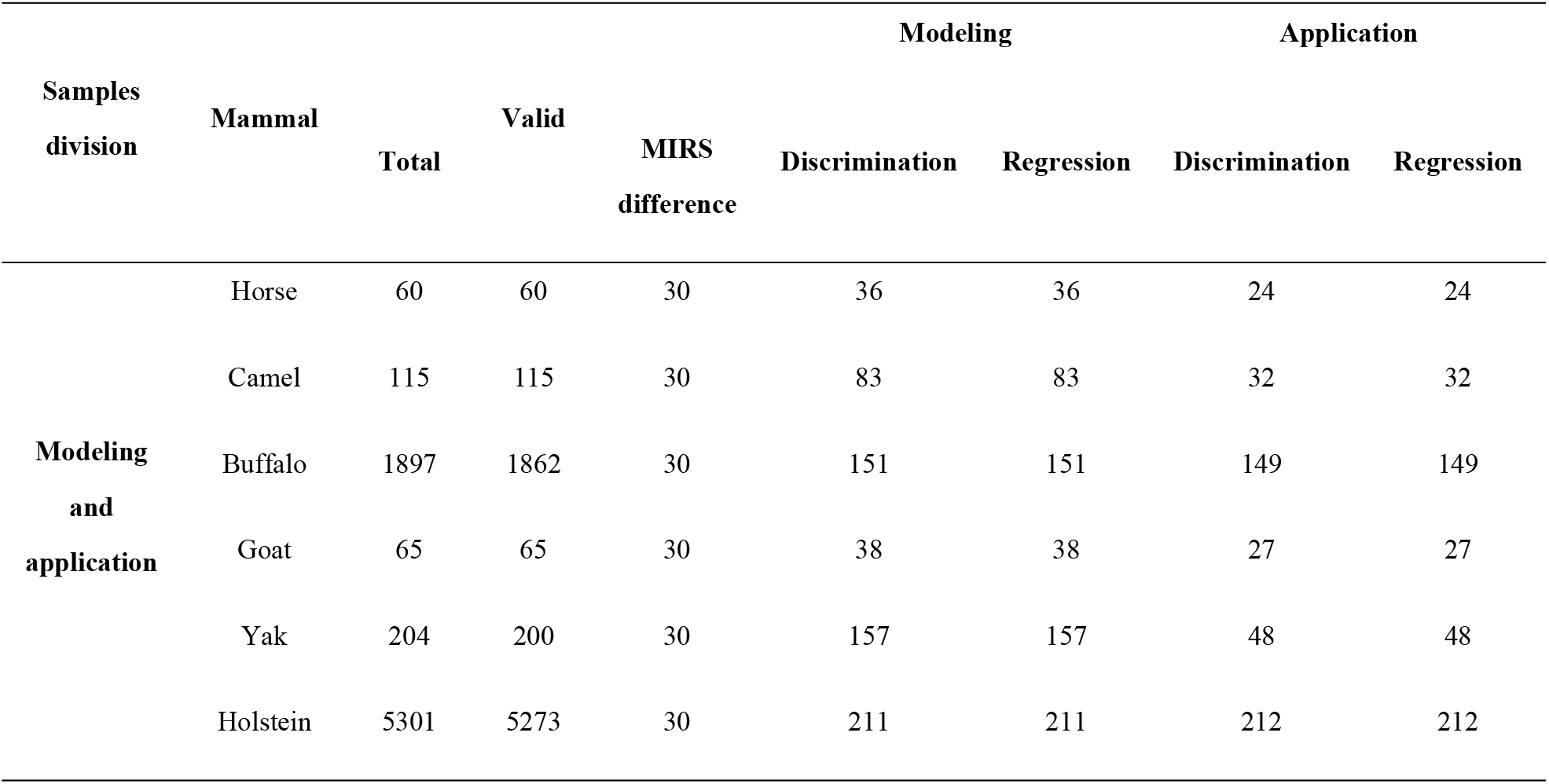
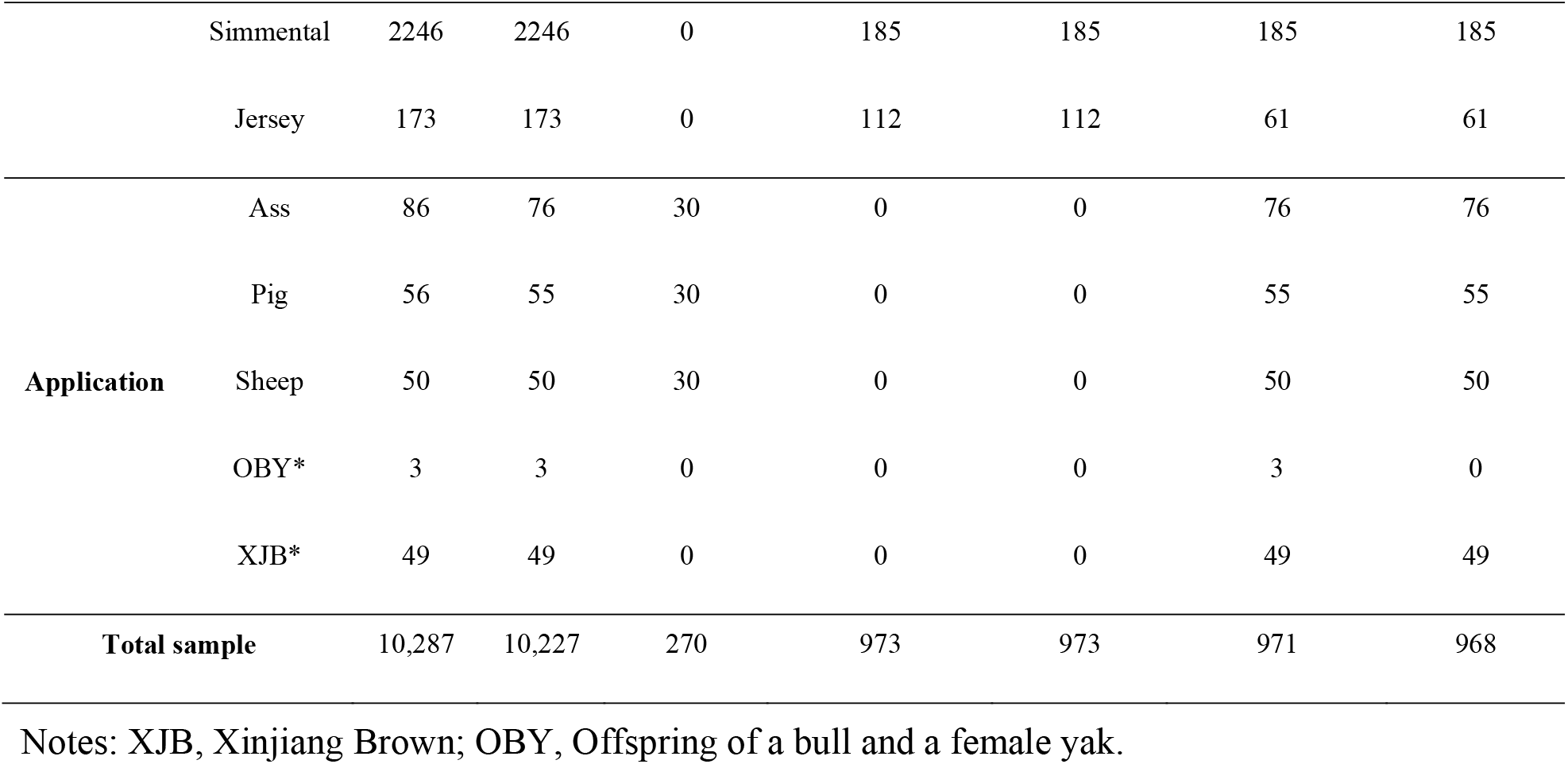
Sample information used in this study.

#### 2.1.2 Differences of MIRS among different animals

The mean spectrum (Fig. 1a) showed the differences in the mean MIRS and the differences in the components of different animal milk samples. The MIRS absorbance of milk fat of buffalo milk and yak milk was higher than that of horse milk and ass milk. The MIRS absorbance of milk protein absorption area of pig milk and sheep milk was higher than that of horse milk and ass milk. The MIRS absorbance of the lactose absorption area of horse milk and ass milk was higher than that of pig milk.

**Fig. 1.**
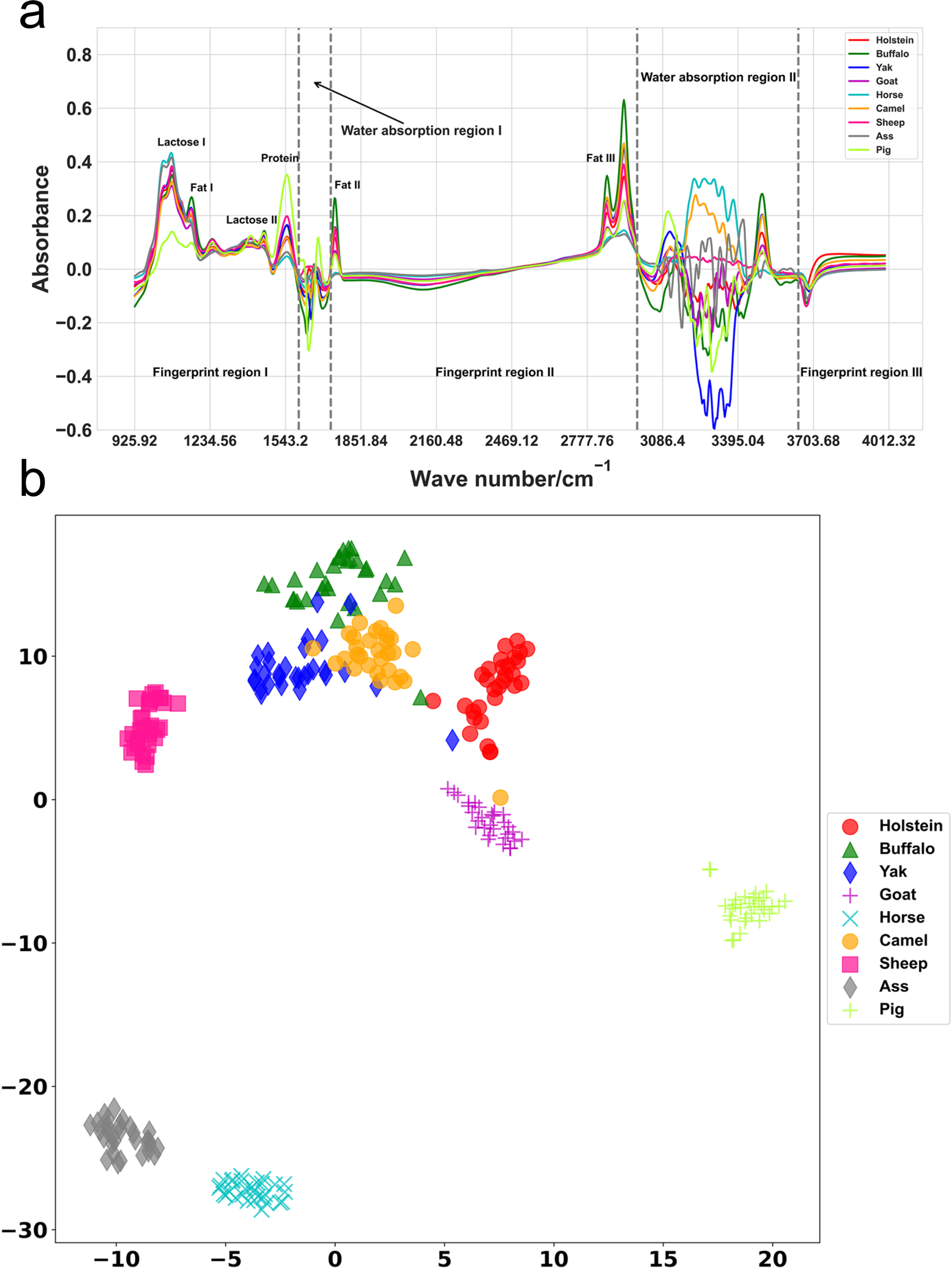
Mean MIRS spectrum and TSNE dimensionality-reduced cluster of 9 animals. **(a)** Mean MIRS spectrum of 9 animals. (b) TSNE dimensionality reduction cluster of MIRS of 9 animals

The 30 samples were randomly selected from different animals with Holstein cows used as the representative of 4 cattle breeds. To reveal the differences in MIRS among 9 different animal species, TSNE (T-distributed stochastic neighbor embedding) dimension reduction and clustering were performed on the MIRS of 9 animal species. As shown in Fig.1b, the clustering effects of all 9 animals were desirable with MIRS of the same species clustered together. Notably, the milk components of the horse, ass, and pig exhibited great differences from those of other species, and they clustered together far away from those of other animals with the best clustering effects. Goat milk and cow milk were clustered close to each other because of their similar components. Our results were similar to the reports by Nicolaou et al. (2010) ^[16]^. The composition of yak milk was similar to that of camel milk and buffalo milk, and their MIRS was also clustered in adjacent positions. Taken together, the MIRS of the 9 animals were quite different which might be attributed to the difference in their milk components. Therefore, MIRS could be used to study the evolutionary relationship of different species of animals.

### 2.2 MIRS-based model construction of animal evolutionary relationships

To explore the use of MIRS in the study of animal evolutionary relationships, the genetic distances (Table S3) of 12 mammals (Table 2, except OBY) were estimated by using whole-genome DNA sequences, and a phylogenetic tree (Fig. S1) was constructed, which provided a reference for establishing animal evolutionary relationships (trees). According to the traditional biological classification, 12 kinds of animals were categorized into Perissodactyla (horse, ass) and Artiodactyla. Artiodactyla was subdivided into Suidae (pig), Camelidae (camel), and Bovidae, and Bovidae was subdivided into Capra (goat), Ovis (sheep), Bubalus (buffalo), and Bos. Bos was divided into Bos grunniens (yak) and Bibos which included Simmental, Holstein, Jersey and Swiss brown (XJB), and other different species. The genetic distance between animals assigned to different evolutionary levels was greater than that between animals assigned to the same level. For example, the genetic distance between pigs and buffalo (0.16229) was greater than that between Jersey and Holstein (0.0018). Additionally, the phylogenetic trees established in this study were similar to those generated by the NCBI Taxonomy Common Tree, indicating the accuracy of the phylogenetic trees of 12 animals constructed in this study, therefore, they could provide the reference for the MIRS-based study of the evolutionary relationship of animals.

To establish the evolutionary relationship (tree) of animals based on MIRS, it is necessary to use MIRS to establish DMEL to determine the branches of the phylogenetic tree reflecting evolutionary level and RMGD to predict the length of the evolutionary tree branches representing genetic distance.

#### 2.2.1 Establishment of DMEL using 8 animal MIRS

In the early stage of this study, 2 algorithms (RF and SVM) and 12 spectral preprocessing methods were compared, and different combinations were formed to establish classification models, respectively. The model established by the 12 preprocessing methods in the combination of the RF algorithm achieved the desired effect, especially, the D1 and D2 preprocessing methods combined with the RF algorithm had a bet classification effect on different animals, and the accuracy and kappa coefficient of the cross-validation sets and test sets were as high as 0.99-1, and thus the combinations of D1+ RF and D2+RF were selected for modeling. The 973 samples from 8 animals were divided into a training set (70%) and a test set (30%) (Table 2), and 5 discrimination models for 5 evolutionary levels were established by using MIRS of 8 animal species. According to the accuracy of the model, a set of optimal models was screened including (1) D1+RF (n_estimators=50, max_depth=8) at order level; (2) D1+RF (n_estimators=50, max_depth=8) at family level; (3) D1+RF (n_estimators=49, max_depth=15) at genus level; (4) D2+RF (n_estimators=50,max_depth =8) at subgenus level; (5) D1+RF (n_estimators=49, max_depth=15)at breeds level). The cross-validation accuracy of these five models was 100%, and the test accuracy was 97% and above (Figure 2a–2e, Table S4), indicating that milk MIRS could be used to establish a very good animal DMEL.

**Fig. 2.**
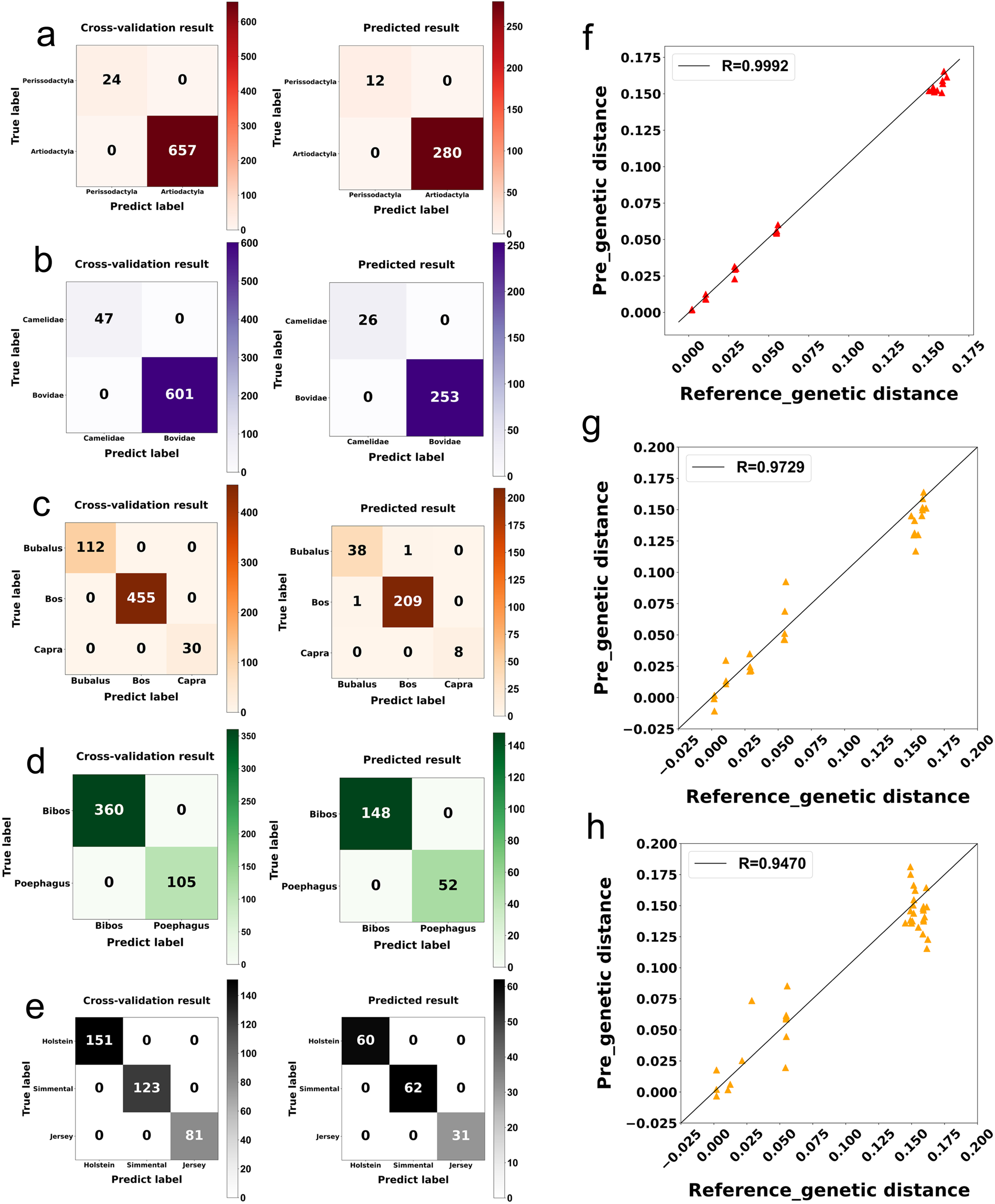
Confusion matrix predicted by five-level DMEL and correlation between genetic distance (GD) predicted based on RMGD and that calculated based on genome-wide. **(a)** The confusion matrix predicted by the order-level model. **(b)** The confusion matrix predicted by the family-level model. **(c)** The confusion matrix predicted by the genus-level model.**(d)** Confusion matrix predicted by the subgenus-level model. **(e)**Confusion matrix predicted by the breeds-level model. **(f)** Correlation between GD of 8 animal species predicted based on the MIR model and that calculated based on the whole genome. **(g)** The correlation between GD of animals that did not participate in the modeling in 8 animal species was predicted based on the MIR model prediction and calculated based on the whole genome. **(h)** Correlation between GD of 4 animal species (for external verification) based on genome-wide calculations and that based on MIR model prediction

#### 2.2.2 Establishment of animal RMGD using 8 animal MIRS

Similar to the establishment of DMEL, the optimal model (D1+SNV+PLSR, n_components=14) was screened from the combination of two D1+SNV preprocessing methods and the PLSR algorithm according to the accuracy of the cross-validation set. Namely, the genetic distance was established by using MIRS of 8 animal species). The correlation coefficient of the cross-validation set of this model was 0.99, and the ratios of GD of the 8 animal species predicted based on the MIRS model to that calculated based on the whole genome were all close to 1 (Fig. 2f, Table S5).

### 2.3 Better validation and application effects of animal evolutionary relationship model established based on MIRS

To verify the effect of the established animal evolutionary relationship model, our established 5 DMELs and 1 RMGD were used to establish the evolutionary relationships of the animals that did not participate in the modeling in 8 species (horse, camel, goat, yak, buffalo, Holstein, Jersey, Simmental) and the animal species that did not participate in the modelings in 5 species (ass, pig, sheep, OBY, and XJB).

#### 2.3.1 Established evolutionary relationships of 8 animals that did not participate in modeling are satisfactory

MIRS and DMEL were used to determine the evolutionary levels of the animals that did not participate in the modeling among 8 animal species (horse, camel, buffalo, goat, yak, Holstein, Simmental, and Jersey). Horses were classified into Perissodactyla by the first-level discriminate model (DM), and camels were classified into Camelidae by the second-level DM. Buffaloes were classified into Bubalus by the third-level DM. Goats were classified into Capra by the third-level DM, and yak was classified into the Bos grunniens by the fourth-level DM. Holstein, Jersey, and Simmental cows were assigned to the corresponding breeds by the fifth-level DM. The population confidence level of the animals that did not participate in the modeling among the 8 animal species was 1. The evolutionary levels of 705 animals (24 horses, 30 camels, 137 buffaloes, 26 goats, 46 yaks, 212 Holsteins, 185 Simmentals, and 61 Jerseys) out of 721 animals were discriminated correctly with the discrimination accuracy rate of 0.98 (Table S6). Therefore, our established five DMELs exhibited excellent discrimination effects at both the population level and individual level of 8 animal species. The evolution levels discriminated based on MIR were the same as those determined based on the whole genome.

Using MIRS and RMGD, we predicted the genetic distance of the 8 animals that did not participate in the modeling with the correlation coefficient of the cross-validation set of 0.97. The genetic distance predicted based on the MIR model was very close to that calculated based on the whole genome (except for yak, Holstein cows, and goats) with the ratio of these two GD values close to 1 (Fig. 2g, Table S7).

#### 2.3.2 Evolutionary relationships established for the five animals that did not participate in the modeling are desirable

MIRS and DMEL were used to judge the evolutionary level of the five animals that did not participate in the modeling (Table S8). The 5 animals (ass, pig, sheep, OBY, and XJB) that did not participate in the modeling were gradually classified into the correct evolutionary levels. The ass was classified into Perissodactyla by the first-level DM. When the pig was classified by the second-level DM, the confidence level was only 0.66, and thus it could not be classified into Camelidae or Bovidae, instead, it was classified into other families. When sheep were classified by the third-level DM, the confidence level was only 0.33, and thus it could not be classified into Bubalus, Capra, or Bos, instead, it was classified into other genera. In addition, the OBY was classified into the Bos grunniens by the fourth-level DM. OBY was a hybrid of bull and female yak, and its evolutionary level was correctly identified, indicating that our model could be used to discriminate the evolutionary level of the hybrid. When XJB was classified by the fifth-level DM, the confidence level was only 0.53, and thus it was classified into other breeds. These results showed that our established five DMELs could be used to effectively discriminate the evolutionary level of animal species including those animal hybrids.

Using MIRS and RMGD, we predicted the genetic distance of 4 animals that did not participate in the modeling (ass, pig, sheep, and XJB) with the correlation coefficient of the cross-validation sets of 0.95. The genetic distance predicted based on the MIR model was highly similar to that calculated based on the whole genome (except for XJB, Jersey, and yak) with their ratio close to 1 (Fig. 2c and Table S9).

Therefore, the animal evolutionary model (including DMEL and RMGD) established in this study exhibited desirable application effect in the establishment of animal evolutionary relationships not only on the animals that did not participate in the modeling among 8 animal species (horse, camel, buffalo, goat, yak, Holstein Simmental, and Jersey, or new unknown animals) but also on 4 animal species that did not participate in the evolutionary relationship modelings (ass, pig, sheep, and XJB, or new unknown animals).

#### 2.3.3 Formulation of application strategies of animal evolutionary relationship model

To better apply the MIRS-based animal evolutionary relationship model, the model application operation flowchart was designed based on the actual applications of the model and analysis of its effect analysis (Fig. 3). The procedures for using the animal evolutionary relationship model were as follows:

**Fig. 3.**
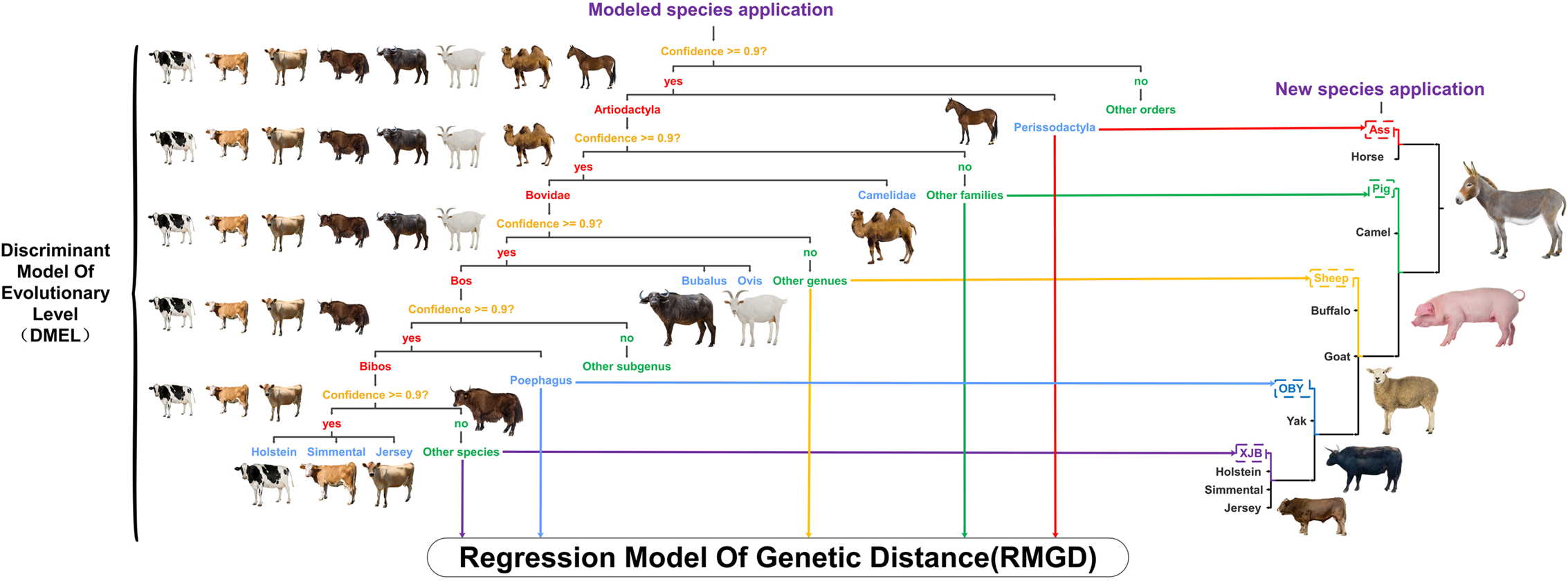
Application flowchart of animal evolutionary relationship model established based on MIRS.

The first step was to use the established 5 DMELs to judge the evolutionary level of animals as follows. (1) The same method of modeling was used to remove abnormal samples from animal milk samples, and 579 MIRS wave points were screened from normal samples; (2) For animal population evolutionary level discrimination, firstly the MIRS mean value of a certain animal was calculated, and the obtained MIRS mean value was introduced as the representative MIRS of this animal species into 5 DMELs for discrimination and classification. After the introduction to the first order level of DMEL, if the confidence level of the animal was 0.9 and above, the animal was classified as Perissodactyla or Artiodactyla. From the second to the fifth level of DMELs, the same level of DMEL confidence level (0.9) was used for discriminating the evolution level of the animals; (3) For animal individual evolutionary level discrimination, all the MIRS values of a certain animal species were substituted into each of 5 DMELs for discrimination and classification. According to the discrimination results of each MIRS in each model, if the discrimination accuracy rate of an animal was higher than 0.9, this animal was determined at this evolutionary level. The discrimination result at the individual level of this animal could be used as the reference for discriminating animal evolution at the population level.

The second step was to use the RMGD to calculate the GD of a certain animal and other animal species as follows. (1) Firstly, abnormal samples were removed, and a total of 278 MIRS wave points of normal samples were screened; (2) The MIRS mean values of this animal and other animal species were calculated, respectively, and the absolute value of MIRS difference between this animal and other animal species was calculated, and 278 MIRS wave points differences between this animal and other animals were obtained. (3) The MIRS wave point difference values were introduced into RMGD to predict the GD between this animal and other animals to clarify the relatedness and evolution degree of this animal and other animals.

If only one or a small number of animal milk samples of a species were available due to the difficulty of obtaining samples, only the animal evolution level discrimination could be performed according to the first step, and the GD prediction in the second step cannot be implemented. In this case, firstly, the MIRS was substituted into DMEL. If the discriminant confidence of the sample at each level was higher than 0.9, the discrimination results of the evolutionary level of this animal were credible. If the confidence level was within 0.8-0.9 at a certain level, the sample had to be subjected to secondary determination. If the confidence level remained stable or at a high level, the discrimination results were credible. If the discrimination confidence level was lower than 0.8, the animal evolution level discrimination results were not credible.

### 2.4 A close relationship among MIRS characteristic waves of different mammalian species, milk components, and their regulatory genes

#### 2.4.1 Close relationship between animal milk MIRS and milk components

Animal milk MIRS was closely related to milk composition, and the large differences in milk components among different animals led to great differences in MIRS (Fig. 1). Perissodactyla and Artiodactyla exhibited large differences in most of the MIRS wave points after removing the water absorption area. For example, the MIRS of horses and asses, both of which belonged to Perissodactyla, had very little difference in absorbance in most bands, but they were quite different from that of other mammals. Camelidae, Suidae, and Bovidae exhibited significantly different absorbances in most bands. The lactose characteristic MIRS absorbance of Suidae was significantly lower than that of Bovidae and Camelidae, and the absorbance of other MIRS regions of Suidae and Camelidae was significantly higher than that of Bovidae. The absorbance of milk fat absorption area of Bubalus was significantly higher. The Bubalus and Ovis displayed significantly different MIRS. Similarly, Bos grunniens and Bibos exhibited significantly different MIRS, and their absorbance in most areas was significantly higher than Bibos. Among the four cattle breeds, the MIRS difference between Jersey and XJB was small, and that between Simmental and Holstein was also small.

The differences in milk components and MIRS of different animals gradually decreased with the increase in evolutionary level, and the differences in milk components and MIRS between Perissodactyla and Artiodactyla (the first level) were significantly higher than those among the four cattle breeds (the fifth level) (Fig. 1, Fig. 4, Table S2). The milk components and MIRS of horses and asses, both of which were Perissodactyla, differed very little, while they differed greatly from other Artiodactyla animals. The MIRS of Camelidae, Suidae, and Bovidae were significantly different. Similarly, Bubalus and Ovis exhibited an obvious difference in MIRS. The MIRS of Bos grunniens was higher than that of Bibos.

**Fig. 4.**
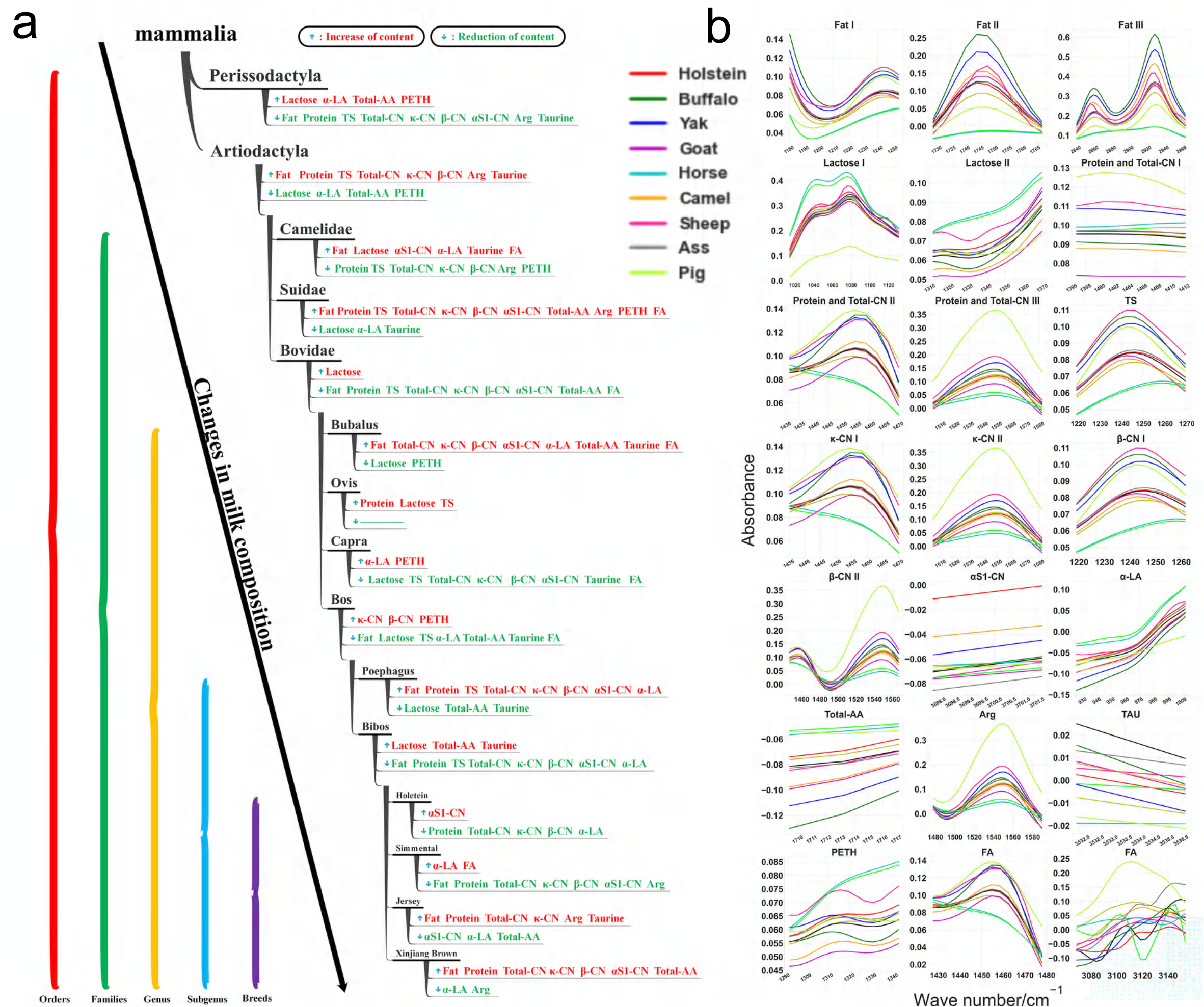
Variation patterns of milk components and their characteristic wavebands during evolution. **(a)** Change patterns of milk components. **(b)** Characteristic wavebands corresponding to different milk components

The MIRS wave points represented by fat, protein, lactose, and others were selected according to the correlation between MIRS wave points in 649.03-3998.59cm^-1^ and milk components (Table 1, Fig. 4), which was consistent with the previous report on the MIRS corresponding to fat, protein, and lactose^[2, 24]^. There was also a correlation between the wave points selected according to the milk components. For example, the MIRS bands selected by protein and total-CN were the same, and the MIRS bands selected by κ-CN and β-CN were similar to those selected by protein. TS and fat exhibited the same bands. These results indicated that the milk components were significantly correlated with MIRS.

In addition, animal DMEL was established based on 23 milk components. It turned out that the discrimination effect of this model was desirable with the kappa coefficient of cross-validation sets and test sets reaching 90% (Table S10), but the discrimination accuracy of the milk component-based model was lower than that of the MIRS-based DMEL (kappa coefficient of >97%), indicating that MIRS was closely related to milk components and that MIRS contained more information on milk components (more than 23 milk components). Therefore, it could be concluded that MIRS could more comprehensively represent the characteristics of different animal milk components and that it was feasible to use MIRS for discriminating animal evolutionary levels.

#### 2.4.2 Milk components corresponding to MIRS characteristic bands are closely related to genes

According to gene function and literature reports, 50 genes related to milk quality were screened. Correlation analysis was carried out between the differences in the average content of milk components corresponding to MIRS characteristic bands of different animals and the genetic distances of 50 genes between different animals (Fig. 5a). The results showed that these genes were closely correlated with milk components. For example, genes *PPP3CA* (R=0.8623) and *PPARD* (R=0.7768) were correlated with lactose; *PRKAA1* (R=0.7820) and *FABP3* (R=0.7439) were correlated with fat; *ACBD6* (R=0.9570) and *SCD* (R=0.9315) were correlated with protein; and *SCD* (R=0.7094)) and *ACBD6* (R=0.6961) were highly correlated with TS. Therefore, the four genes (*PPP3CA, PRKAA1, ACBD6*. and *SCD*) with the highest correlation with the four milk components were selected, and the four genes from different animals were subjected to multiple sequence alignment to construct A phylogenetic tree. Figure 5b showed that for most mammals, single gene alignment-based phylogenetic trees were similar to whole-genome sequence alignment-based ones. However, there were also a few exceptions, for example, the *ACBD6, PPP3CA*, and *SCD* genes of pigs were clustered separately, but the *PRKAA1* gene was clustered with the camel. The possible reason for these exceptions might be that pig colostrum and normal milk had higher milk protein, higher milk fat, total solid contents, and lower lactose content than those of other animals, further indicating that these genes regulated the corresponding milk components.

**Fig. 5.**
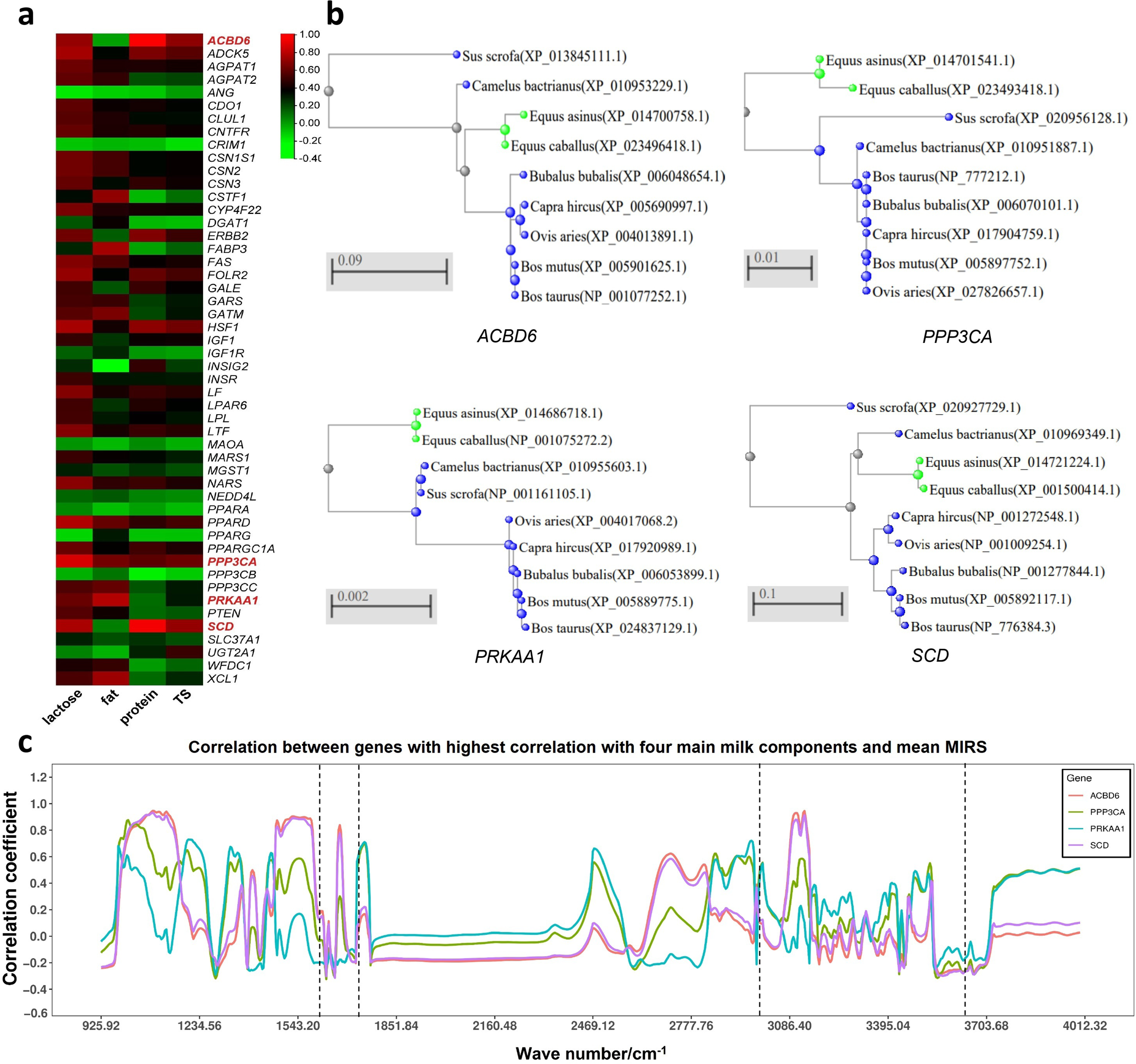
Correlation between 50 genes and 4 main milk components. **(a)**Heat map of correlation between 50 genes and four main milk components. **(b)**Cluster analysis of genes with the highest correlation with four main milk components. **(c)** Correlation between genes with the highest correlation with four main milk components and mean MIRS.

#### 2.4.3 Animal MIRS is closely related to genes

To further analyze the relationship between MIRS characteristic waves and genes, the correlation analysis was carried out between the differences in the genetic distances of the above-mentioned four genes (*PPP3CA, PRKAA1, ACBD6*, and *SCD*) among different animals and the differences in MIRS characteristic waves of different animals. Figure 5c showed that the MIRS characteristic waves with the highest correlation with the gene *ACBD6* were 1087.96-1091.81 cm^-1^, 1477.61-1516.19cm^-1^, and 3097.97-3132.70 cm^-1^; the MIRS characteristic waves with the highest correlation with *PPP3CA* were 1003.08-1060.95 cm^-1^, 1739.96-1743.82 cm^-1^, and 2847.20-2966.80 cm^-1^; the MIRS characteristic waves with the highest correlation with *PRKAA1* were 1192.12-1222.99 cm^-1^, 1743.82-1759.25 cm^-1^ and 2959.09-2974.52 cm^-1^; those with the highest correlation with *SCD* were 1057.09-1057.09 cm^-1^, 1512.34-1581.78 cm^-1^, and 1581.78-3136.55 cm^-1^.

The MIRS characteristic waves with high gene correlation (R>0.6) were compared with MIRS characteristic waves representing different milk components (screened based on the correlation between MIR characteristic waves and milk components). The results showed that most of the MIRS characteristic waves screened by these two methods were overlapped, such as lactose LL absorption areas Lactose I and Lactose II, and milk fat LF absorption areas Fat I, Fat II, and Fat III, and others. (Fig. 1 and Fig. 5c). The genes *ACBD6* and *SCD* showed a high correlation even in the water absorption area. In addition, more bands were screened based on the correlation between genes and MIR characteristic waves, which also included bands that could not be screened out according to the correlation between milk components and MIR characteristic waves (such as 2469.12-2476.84 cm^-1^).

Therefore, the MIR characteristic waves of different animals were closely related to genes, and these correlations are mediated by specific substances in the milk components. The differences between the genes of different animals were reflected in the differences in the genetic distances of different genes, and thus the changes in genetic distances tended to result in the change in milk composition. The changes in milk composition were finally reflected by the MIR characteristic waves, and thus animal milk composition, animal MIRS, and animal genes were closely related (Fig. S2). MIRS could indirectly reflect gene differences and their genetic distances. Therefore, it was possible to use MIRS to establish the evolutionary relationship of mammals.

## Discussion

### A new method for discriminating animal evolutionary relationships based on MIRS and its application effects

This study associated mammalian milk MIRS with animal evolution for the first time, and established a novel method for analyzing animal evolutionary relationship models (including 5 DMELs and 1 RMGD) based on MIRS and machine learning. It broke through the traditional animal evolution research pathway which mainly relied on animal gene nucleotide sequence technology ^[25–28]^.

This study employed the established MIR-based animal evolutionary relationship model to discriminate the evolutionary levels and estimate the genetic distance of 12 different species of animals (including those which did not participate in the modeling). The results showed that our model had ideal application effects in establishing animal evolutionary relationships at five levels of order, family, genus, subgenus, and species, in revealing genetic distance, and in discriminating the genetic relationships of animal individuals and populations. In our established evolutionary level identification model, MIRS was used to replace the whole genome for discriminating the animal evolution levels with an accuracy of up to 98%. Although our established RMGD could not completely replace the whole genome with MIRS to calculate the animal genetic distance, the genetic relationship of animals could be preliminarily determined according to the predicted genetic distance. The ratio of the GD predicted by our model to that obtained by the whole genome was close to 1.

To make better use of our model, we formulated the protocols and strategies for using animal evolutionary relationship models. In DMEL, confidence was used as the main basis for confirming the evolutionary level of a certain animal. For example, among 5 different species of animals (ass, pig, sheep, OBY, and XJB), ass belonged to the existing branch in the model with its confidence level reaching 0.9, and the confidence level of OBY also reached 0.9. However, pigs, sheep, and XJB cattle were the branches that did not exist in the model, and all their confidence levels were 0.7, and thus the confidence level could be used as a direct criterion in a model to determine whether an animal belonged to an existing branch in the model or not. As for the application of RMGD, if the genetic distance (GD) predicted by the model matched the determined animal evolution level, the results predicted by RMGD were reliable since the genetic distance was to discriminate the evolutionary differences between animal species from a quantitative point of view.

### Evidence and feasibility of establishing MIRS-based animal evolutionary relationship model

MIRS is the fundamental absorption band of organic matter and inorganic ions, and it is most suitable for qualitative and quantitative analysis of substances^[2]^. When the contraction and bending vibration frequencies of specific chemical bonds of molecular functional groups in milk are consistent with the MIRS frequency, the molecular absorption energy rises from the ground-state vibrational energy level to the vibrational energy level with higher energy, and the mid-infrared light of this wavelength is absorbed by the substance, thus resulting in a light transmittance value, which is finally manifested as a specific mid-infrared absorption peak. This peak can be used to analyze the presence or absence and the content of a certain component in milk^[24, 29]^. Animal milk MIRS has been widely used in milk quality testing and DHI^[30]^.

The DNA sequence of animal genes may change during evolution^[31]^, which also affects the regulation of mammalian milk production and milk components, which is the main reason for the differences in milk components and corresponding MIRS among different animal species^[32]^. Animal milk MIRS can comprehensively reflect MIRS fingerprints of milk components, and can also be used as phenotypic data for milk quality traits^[33]^. In this study, five DMELs (order, family, genus, subgenus, breeds) established based on 23 milk components from 12 mammal species were used to predict the evolutionary level of animals. The results showed that the milk components of different animal species were quite different, that the correlation between MIR characteristic waves of different animals and genes was mediated by specific milk components, and that the genetic differences of different animals were reflected by the difference in genetic distance of different genes, that the change of genetic distance of genes led to the change of milk components, and that the change of milk components was finally reflected by the MIR characteristic waves. Therefore, MIRS could be used to indirectly reflect the difference in genes and their genetic distance, which was supported by one previous study using NIRS (Near-Infrared Spectroscopy) for the investigation of the genetic distance of plants ^[34,35]^. These findings preliminarily clarified the scientificity and feasibility of using MIRS for studying the evolutionary relationship of animals.

### MIRS-based animal evolutionary relationship model remains to be improved

Due to the limited number of animal species (12 species) used in this study, the small sample size, and the limited distribution of samples, the MIRS feature information learned by the model may be limited. In addition, preliminary experiments in our laboratory showed that the established evolutionary relationship model had similar application effects in the colostrum and normal milk of Holstein cows, and thus the colostrum of pigs (with the milk of other animals used as normal milk) was collected as experiment materials to further explore the application effect of the model in colostrum. However, due to its high protein content and high dry matter content, pig colostrum needed to be diluted at a ratio of 1:1 before measurement. Furthermore, sheep MIRS was measured by BENTLEY milk composition analyzer, while MIRS of other animal milk samples was measured by FOSS milk composition analyzer. Since the band ranges of MIRS measurement of these two instruments were different, their common bands were selected for research. All these factors may have a certain impact on the research results^[36]^, and thus it is necessary to collect more samples with high representativeness and diversity to further improve the model to increase the prediction ability and universality of the model.

Considering that a large number of mammalian milk samples can be collected non-invasively and conveniently, and MIRS detection technology has multiple advantages such as rapidity, high efficiency, and low cost, it is necessary to develop new technologies for MIRS-based animal evolution and genetic correlation research.

## Conclusion

In this study, the relationships among the animal milk MIRS characteristic waves, the corresponding specific milk components, and regulatory genes were analyzed, which provided empirical evidence and a theoretical basis for establishing a mammalian evolutionary relationship based on animal milk MIRS. Based on animal milk MIRS, a mammalian evolutionary relationship model was further constructed including 5 animal DMELs and 1 animal RMGD. Within the scope of the test samples, the accurate evolutionary relationship among different mammals was established, and a good model application effect was achieved. However, our established models remain to be further improved and optimized by applying them to more samples with high representativeness and diversity to improve the accuracy and universality of the model. Our data reveal that animal milk MIRS can replace the genome to evaluate the evolutionary level of mammals. Although MIRS can not completely replace the whole genome to calculate the genetic distance of animals, it can preliminarily predict the genetic distance of animals, thus estimating the genetic relationship. Since MIRS has multiple advantages such as rapidity, high efficiency, low cost, accuracy, and non-invasiveness, it is essential to develop a new animal milk MIRS-based technology for research on the animal evolutionary relationship. Overall, this study provides new approaches and perspectives for mammalian evolution and genetic research.

## Materials and Methods

### 4.1 Collection of test materials

Normal milk of twelve animal milk samples and pig colostrum (Table 2) were collected from 15 provinces and regions in China, and a total of 10,287 milk samples were obtained. Following DHI milk collection and measurement technical regulations, 30-50ml milk was collected using a flow meter or manually and mixed evenly during the whole milking process. A small amount of Bronopol preservative was added to the collected milk sample to dissolve and mix evenly with the milk sample. The milk samples were quickly sent to the DHI laboratory at 2-4°C.

### 4.2 Determination of 4 main components and MIRS, data collection, and prediction of 19 milk component contents

The Bentley.NexGen-400 milk composition analyzer (Minnesota, USA) was employed for the determination of sheep milk. The FOSS Milkoscan FT + milk composition analyzer (Hilleroed, Denmark) was used for the milk component determination of 13 types of animals. Due to the high dry matter content of pig colostrum, the determination was performed after 1: 1 dilution with water and the measured values of 4 main milk components was multiplied by 2. Four main milk components (milk fat, milk protein, milk lactose, and total solid (TS)) and MIRS data were collected.

We predicted 19 milk component contents of different animal milk samples including 5 proteins (total casein, κ-CN, β-CN, αS1-CN, and α-LA, in g/L), 4 amino acids (total free amino acids, arginine, taurine, and phosphoethanolamine, nmol/L), and 10 fatty acids (total fatty Acids, saturated fatty acids, monounsaturated fatty acids, polyunsaturated fatty acids, C12:0, C14:0, C16:0, C18:0, C18:1, and C18:2, g/100g) using the milk MIRS-based prediction model (R>0.8, Table S1) established in our laboratory earlier work. Since the predicted values of three main component contents (protein percentage (PP), fat percentage (FP), total solid (TS)) by buffalo milk MIRS-based model (R>0.95) established in our laboratory earlier work (unpublished) were not significantly different from the measured values by cow milk model-based DHI detection method or the reference values measured by chemical method (P>0.05), the cow milk model could be used for predicting the contents of other animal milk components to a certain extent.

We eliminated 60 abnormal samples with missing or incomplete MIRS data, MIRS-based Mahalanobis distance≥ 3, four major milk component contents below 0% by DHI detection, FP>25%, PP>19%, LP>7.5%, and TS>38%. After elimination, a total of 10,227 valid milk samples were screened from 10,287 milk samples collected from 13 animals (Table 1; OBY has 3 milk samples).

### 4.3 Construction of animal evolutionary tree based on the whole genome

To effectively reflect the evolutionary relationship of animals, the whole genome sequences were used to construct the phylogenetic tree. The reference genomes of 12 animals (except OBY) were selected from NCBI, including Pig (GCF_000003025.6_Sscrofa11.1), Horse (GCF_002863925.1_EquCab3.0), Ass (GCF_016077325.2_ASM1607732v2), Camel (GCF_000767855555555) 0), Goat (GCA_001704415.1_ARS1), Sheep (GCA_016772045.1_ARS-UI_Ramb_v2.0), Yak (GCA_005887515.2_BosGru3.0), Buffalo (GCA_003121395.1_UOA_WB_1), and 4 cattle breeds Holstein (GCA_021347905.1_ARS-LIC_NZ_Holstein-Friesian_1), Jersey (GCA_021234555.1_ARS-LIC_NZ_Jersey), Simmental (GCA_018282465.1_ARS_Simm1.0), and Xinjiang brown cattle (XJB, since no XJB reference genome was available, brown Swiss cattle with relatively closest kinship (GCA_914753205.1_Brown_Swiss_cow) was selected as substitute). The genetic distances of 12 animals were estimated by running MASH software with default parameters^[37]^, and the obtained genetic distances were submitted to ITOL (Interactive Tree Of Life) to construct an evolutionary tree^[38]^.

### 4.4 Method for constructing animal evolutionary relationship model using MIRS

The branch (evolutionary level) and branch length (genetic distance) in the animal phylogenetic tree (reflecting evolutionary relationship) were the main indicators of the evolutionary relationship of animals. Therefore, MIRS was used to establish the discriminant model of evolutionary level (DMEL) and regression model of genetic distance (RMGD), respectively, and then based on these two models, animal evolutionary level and genetic distance were obtained to further establish the animal evolutionary relationship. A total of 973 samples (Table 1) from 8 animals (Holstein, Simmental, Jersey, yak, goat, buffalo, camel, and horse) were used for modeling.

#### DMEL

According to the phylogenetic tree constructed based on the whole genome and the determined animal evolution level, A total of 5 DMELs of the 5 evolutionary levels (order, family, genus, subgenus, species) (5 in total) were established using MIRS (each DMEL)for one level. The specific steps were as follows: (1) MIRS (925.92-1543.2cm^-1^, 1720.67-2970.66cm^-1^, and 3634.24-3998.59cm^-1^) which could be determined by both FOSS (925.92-5011.54cm-1, 1060 wave points) and Bentley (649.03-3998.59cm-1, 899 wave points) milk composition analyzers were selected to obtain 579 MIRS wave points for modeling; (2) The classification models were established based on two machine learning classification algorithms (random forest (RF) and support vector machine (SVM)) and the different combinations of 12 MIRS preprocessing methods (no preprocessing (None), normalization (MMS), standardization (SS), Zero-centered CT, standard normal variable (SNV), moving average (MA), smoothing filter Savitzky Golay SG, and multiplicative scatter correction (MSC), first difference method D1, second order difference D2, detrend correction (DT), wavelet transform (WAVE) to screen the optimal combination of machine learning algorithms and preprocessing methods, respectively; (3) Five animal DMELs for each of 5 evolutionary levels were established based on 579 MIRS wave points from some (973) samples of 8 animals and the optimal combination of preprocessing methods and algorithms, respectively; (4) The model effects were compared, and they were further compared with the whole-genome analysis results to screen one set of optimal animal DMELs (composed of 5 models).

#### RMGD

Animal RMGD was established using the mean MIRS of each animal and animal genetic distance calculated based on the whole genome. The steps were as follows: (1) MIRS average values of 973 samples from 8 animals were calculated, and then the absolute values of the differences in MIRS average value between animals were calculated. A total of 28 MIRS difference values were obtained; (2) To establish the best regression model, 327 MIRS wave points were preliminarily screened from 579 wave points using 5 wave point selection algorithms (GA, UVE, SPA, Lars, and Cars) with 269, 1, 8, 40, and 54 screened from these 5 algorithms, respectively; (3) Using the genetic distance calculated based on the whole genome as the reference value, the modeling was performed on 28 MIRS difference values with 327 MIRS points based on different combinations formed from 12 preprocessing methods (None, MMS, SS, CT, SNV, MA, SG, MSC, D1, D2, DT, and WAVE) by partial least squares regression (PLSR) algorithm. The optimal combination of preprocessing method and PLSR algorithm was screened. Further modeling was performed by the PLSR algorithm, and 278 MIR wave points were screened from 327 MIRS wave points by manual selection method to obtain optimal dimensionality reduction parameters; (4) The different model effects were compared, and further model effects and the whole genome prediction results were compared to screen the optimal RMGD.

### 4.5 Analysis of relationship among animal milk characteristic wavebands, milk components, and genes

#### Method 1

Firstly, we selected the milk components such as lactose, milk fat, and milk protein represented by animal MIR characteristic wavebands, and calculated the difference in the average content of a certain component among various animals. Secondly, combining literature reports, we selected important genes related to this component from the NCBI Orthologs gene database and submitted these gene sequences to NCBI’s multiple sequence alignment tool COBALT (Constraint-based Multiple Alignment Tool) for alignment. Thirdly, we imported the alignment results (.aln) into MEGA11for processing with default parameters and calculated animal genetic distance based on gene sequences. Finally, correlation analysis was performed between the differences in average content of milk components and the gene sequence-based genetic distances of animals, and the results were visualized in the form of heat maps.

#### Method 2

The difference in the MIR characteristic waves among different animals was directly selected, and the correlation between these MIR characteristic wave differences and the whole genome-based animal genetic distances was analyzed with the results visualized using ggplot2 in the R package.

## Acknowledgements

Financial assistance from the Inter-Governmental International Science and Technology Cooperation Project of the State Key Research and Development Program (2021YFE0115500) and International Science and Technology Cooperation Project of Hubei Province. We are also grateful to Hubei Dairy Herd Improvement Center, Henan Dairy Herd Improvement Center and Hebei Province Animal Husbandry and Improved Breeds Work Station for testing and providing the data of animal milk used in this study, and Prof. Ping Liu of college of foreign languages of Huazhong Agricultural University for her efforts and kindly suggestions in the writing and correcting this manuscript.

## Conflicts of Interest

The authors declare no competing interests.

## Data availability statement

The datasets generated during and/or analysed during the current study are not publicly available due to a data confidentiality agreement with the partner unit was signed but are available from the corresponding author on reasonable request.

